# Accretion of novel genetic elements reverses genome erosion and enables host-switching in the deep sea

**DOI:** 10.64898/2026.10.22.761039

**Authors:** Camryn N Morey, Russell Corbett-Detig, Shelbi L Russell

## Abstract

Endosymbiont genome erosion is driven by relaxed selection and isolation from free-living relatives’ gene pools, and can be stalled by horizontal transmission events, but it remains unknown whether genome erosion can be reversed. The chemosynthetic gammaproteobacterial symbionts of deep-sea clams in the family Vesicomyidae offer a useful test case for whether eroded symbiont genomes can acquire novel functional content. Despite having some of the smallest genomes among marine endosymbionts, several lines of evidence, including divergent 16S rRNA variants within hosts, host-switching in sympatry, and phylogenetic signatures of repeated host transitions, suggest these symbionts retain more environmental exposure and evolutionary flexibility than expected under strict vertical transmission. Here, we use comparative genomics across the vesicomyid symbionts to test for evidence of functional gene gain. We find that three lineages have each independently expanded their genomes by approximately 25% relative to close relatives, with some expansions driven by insertion sequence (IS) or phage acquisition. The genomes that contain IS elements have disrupted genome synteny genome-wide, suggesting that these elements mediated structural rearrangements through ectopic recombination. Most strikingly, the symbiont of a thyasirid clam host, *Thyasira* sp. Haima SOB, harbors four unusually large (∼24–40 kb), highly repetitive genes with structural homology to adhesins found broadly across prokaryotes and eukaryotes. These genes show hallmarks of horizontal acquisition and IS-mediated mobilization, and their predicted export via a Type I secretion system suggests a role in mediating attachment to, or exchange of metabolites across, host cell surfaces. Considering that this symbiont lineage switched from a vesicomyid host to a thyasirid host relatively recently, and thyasirids typically host extracellular symbionts, these adhesins may have been instrumental in this host-switch. Together, these findings provide evidence that genome erosion in intracellular symbionts is not strictly unidirectional, and mobile genetic elements can serve as vectors for functional innovation, potentially enabling adaptation to new hosts and environments even after substantial genome reduction has occurred.

## Introduction

Intracellular symbiotic bacteria typically have small, gene-dense genomes compared to their free-living relatives due to reductive genome evolution (RGE) [1]. Upon the establishment of symbiosis and host restriction, small population sizes and transmission bottlenecks reduce recombination, weakening the efficacy of selection against new mutations, therefore driving genetic drift and loss in regions of the genome no longer necessary for life inside of host cells (Figure 1). Nonetheless, the paradigm of loss and reduction does not explain patterns in genome size seen in some ancient host-symbiont associations. Many lineages of ancient symbionts in marine environments do not have genomes as small as expected, considering the evolutionary age of these relationships [2–5]. Occasional horizontal transmission and recombination has been shown to “reset” the reductive evolutionary trajectory by reintroducing genomic variation, even in nearly strictly vertical transmitted symbionts [4, 6].

**Figure 1.**
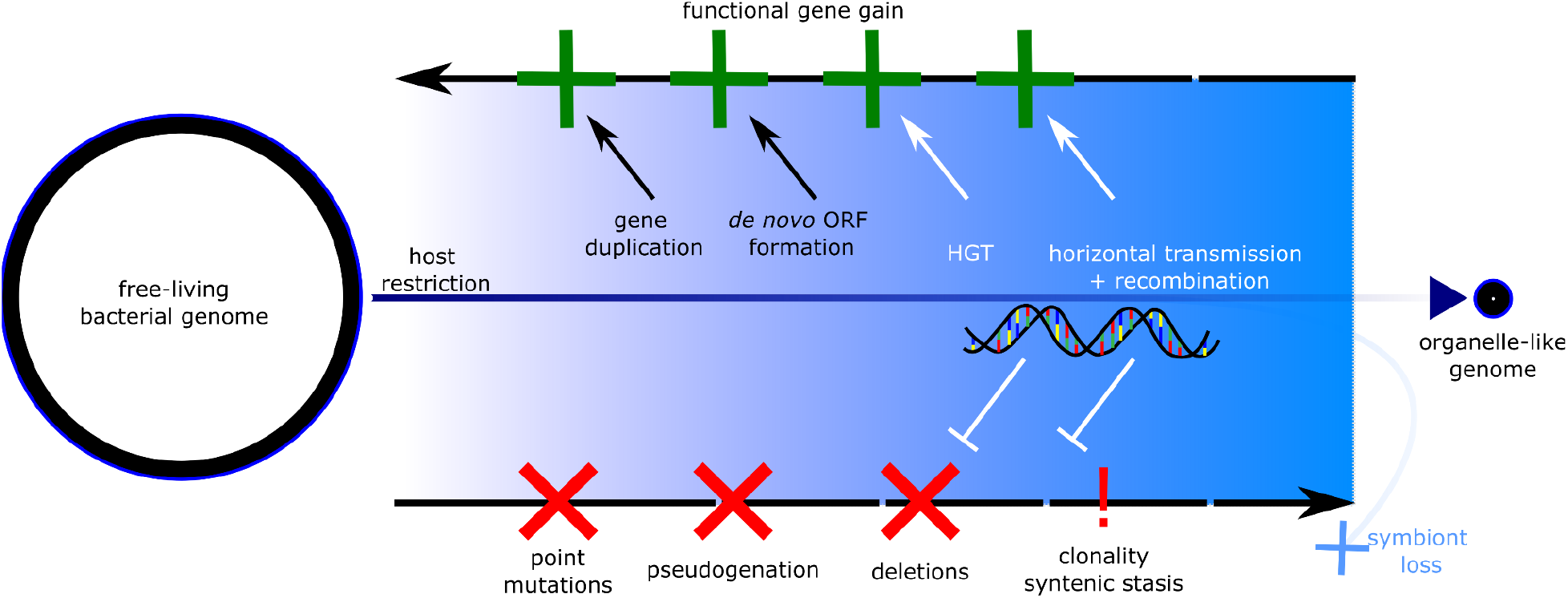
Genome erosion may be reversible. Here, we explore whether genome erosion can be reversed by functional gene gain, in addition to being stalled by HGT and homologous recombination. New genes may arise by HGT, as well as gene duplication and de novo ORF formation, as occurs in eukaryotes.

While occasional recombination can stall erosion by introducing and maintaining genetic variability, it remains unexplored to what extent symbiont genome function can be *gained* despite the overall trend towards loss. In addition to rescuing loss of function alleles, HGT and homologous recombination can introduce functional genetic content that could be leveraged in symbiosis [7–10]. Whether MGEs contribute to gain in ancient symbionts is unclear, as they are largely absent from strictly vertically transmitted lineages, where drift and the cost of proliferation in an already reduced genome favor their loss.

Chemosynthetic symbionts supply their hosts with nutrition through sulfur or methane oxidation in ecosystems with little photosynthetically derived organic carbon [11, 12]. They have been acquired independently by many marine invertebrates [13] and vary widely in transmission mode and genome size [14–16], making them a useful system for exploring the dynamics of gene gain and loss in symbionts over long evolutionary time scales. The gammaproteobacterial symbionts associated with deep sea clams in the family Vesicomyidae may still be capable of acquiring new functions through gene gain, despite having some of the smallest genomes among marine endosymbionts. Although all known vesicomyid symbionts are intracellular and vertically transmitted [17, 18], evidence suggests they have ample exposure to the environment: divergent 16S rRNA sequences have been detected in some host individuals [19, 20], some symbionts have been reported to jump between hosts in sympatry [21], and phylogenetic reconstructions indicate many host-switching events occurred in their evolutionary history [22].

Consistent with the idea that the vesicomyids symbionts retain the ability to switch hosts, a bacterium that clusters within the *Ruthia* clade of vesicomyid symbionts, referred to as Clade II, was recently found in a thyasirid clam from the Haima cold seep in the South China Sea [23]. It and two other Clade II symbionts have genomes ∼300 Kb larger than the largest previously sequenced vesicomyid symbiont genomes [24, 25], suggesting either retention of ancestral genome content or recent expansions resulting from the acquisition of novel genetic material. Additionally, symbionts of thyasirid clams differ from those of vesicomyid clams in that they are typically located extracellularly on the gill epithelium [26]. For an intracellular symbiont to colonize a new bivalve species that normally hosts extracellular symbionts, it potentially requires novel functions to aid in intracellular colonization or the extracellular exchange of metabolites, which could be encoded in these putatively expanded regions.

Here, we assemble a comparative genomics dataset for the vesicomyid and vesicomyid-like symbionts and test for evidence of independent genome expansions and functional gene gain over their histories. We find that previously unreported mobile elements dominate the three larger Clade II symbiont genomes, with likely origins after divergence from their common ancestor, indicating larger genomes are indicative of recent expansions rather than patterns of differential loss. We also find the presence of potentially novel coding sequences in these genomes, including large adhesins in a host-switching symbiont, which might have mediated host adaptation during horizontal transmission events. These results build upon existing evidence that reductive genome evolution in symbionts is not linear, and infrequent genetic exchange between symbiont populations and extracellular microbes can act as a source for functional novelty even in ancient obligate symbiont lineages.

## Results and Discussion

### Clade II vesicomyid symbionts have experienced at least three independent genome expansion events

Three of the Clade II symbionts have genomes approximately 257–351 kb (XXX%---XXX%) larger than the average genome size of other Clade II relatives. These “extra” genomic regions contain a large number of mobile elements which are nearly or completely absent from their closest relatives (Figure 2A). The mobile element type and diversity is variable between the three genomes. Insertion sequences dominate the genomes of the *Isorropodon fossajaponica* SOB and *Thyasira* sp. Haima SOB genomes, contributing to 6.51% and 6.94% of the genome size, respectively (Supplementary Table S2). The *Isorropodon fossajaponica* SOB genome contains mostly IS3-family insertion sequences, with 86 copies, as well as 3 IS30-family elements and 1 IS256-family element (Figure 2B). The closest relative of *Isorropodon fossajaponica* SOB, *Pliocardia ponderosa* SOB, contains one copy of an IS3-family element, but does not share any other elements. The *Thyasira* sp. Haima SOB genome contains a greater diversity of IS elements, but mostly consists of IS3-family and IS5-family insertion sequences, with 43 IS3-family copies and 37 IS5-family copies, along with five IS1595-family copies, five IS256-family copies, three IS30-family copies, and three IS481-family copies (Figure 2C). The high copy number of insertion sequences across these independent lineages suggests recent acquisitions and proliferation in these three genomes, resulting in an expansion in genome size.

**Figure 2.**
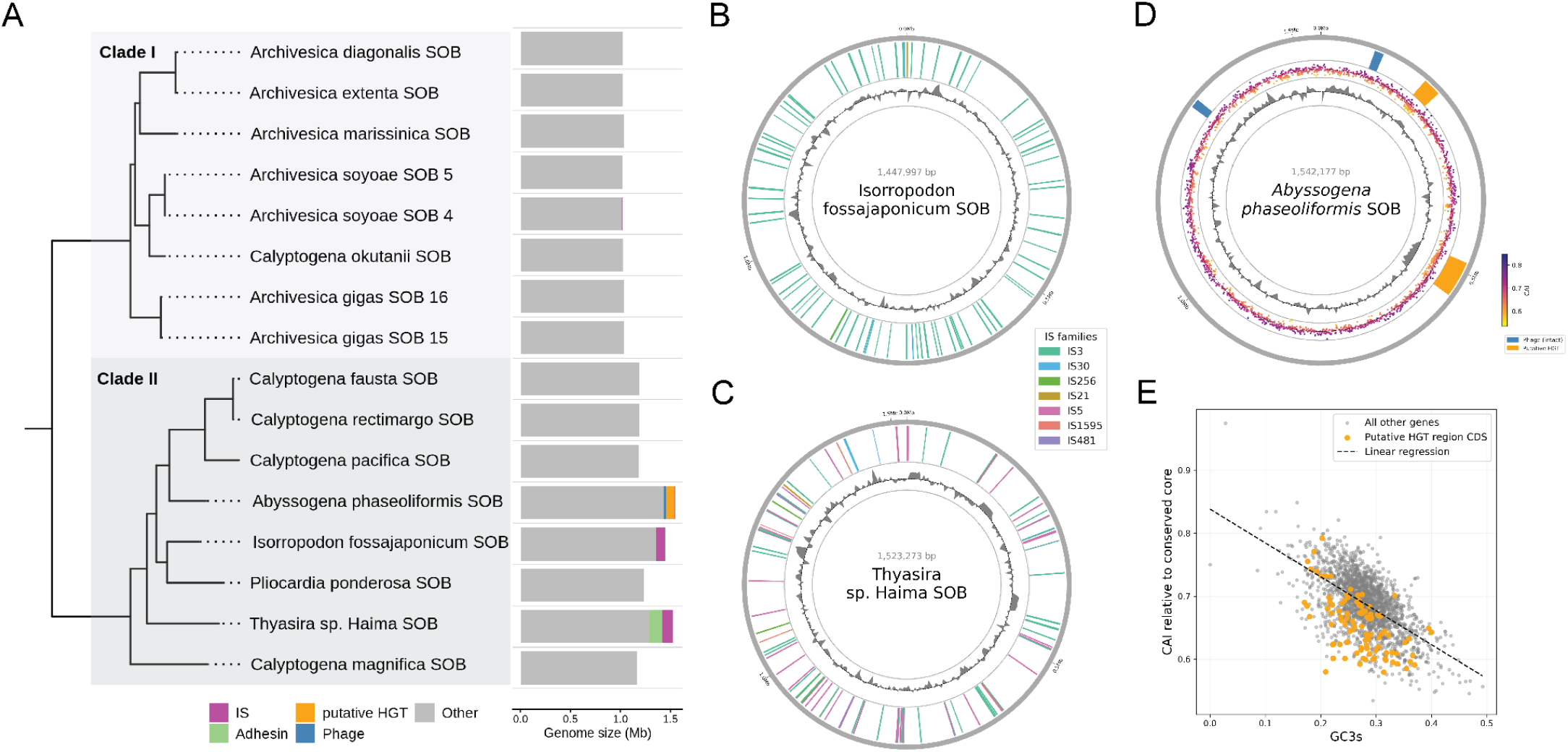
Genome size variation and mobile element distribution across vesicomyid and vesicomyid-like symbionts. (A) Maximum likelihood phylogeny based on single-copy orthologs (left) and genome size variation and mobile element composition across Clade I and Clade II symbionts (right). Insertion sequence distribution (outer ring) and GC content (inner ring) across the genomes of (B) *Isorropodon fossajaponica* SOB and (C) *Thyasira* sp. Haima SOB, colored by predicted IS family. (D) *Abyssogena phaseoliformis* SOB phage and putative HGT region (outer ring) distribution, codon adaptation index (CAI) of coding sequences by start site (middle ring), and GC content (inner ring). (E) Correlation between CAI and GC content at the third codon position (GC3) for all coding sequences in the *Abyssogena phaseoliformis* SOB genome, with orange markers for coding sequences occurring in putative HGT regions.

While insertion sequences are absent from the *Abyssogena phaseoliformis* SOB, the presence of other mobile elements indicates recent acquisition from currently unsequenced origins (Figure 2D). Two 10,817 bp intact prophages are present, which encode viral proteins and have no similarity to any currently sequenced virus or organism, while prophages and any related fragments are absent from all other Clade I and Clade II relatives. Additionally, the *Abyssogena phaseoliformis* SOB genome contains two large, putative HGT regions of lengths 56,250 bp and 27,112 bp. If these regions were a result of ancestral retention, we would expect homologs or fragments of related sequences to be present in symbiotic or free-living relatives within Thioglobaceae. However, genes in these regions have no detectable homologs in relatives, nor do they have similarity to any currently sequenced organism (Supplementary Figure S1). Furthermore, these regions have a notably different GC content, and contain coding sequences that deviate in codon usage (p < 0.05 for both regions, independently) compared to core genes conserved across Clade I and Clade II (Figure 2D), even when accounting for the effect of GC content on codon usage (Figure 2E).

### Clade II vesicomyid symbionts experienced several IS element invasions from environmental sources

Heterogeneous IS element composition among the vesicomyid genomes reveals ancient and more recent transfers from non-ancestral DNA sources (Figure 3 and Supplementary Figure S2). Although the IS3 element was detected in both *I. fossajaponica* SOB and *Thyasira sp*. Haima SOBs’ genomes, many of the copies were more closely related to IS3 elements found in other host-associated metagenomes, such as sponges, annelids, and clams in other families, or elements from environmental metagenomes in the TARA Oceans collection [27] (Figure 3A). This suggests that the IS3 elements in these genomes arose from multiple independent novel insertions and subsequent expansions across these genomes. The IS5 element was only detected in the *Thyasira* sp. Haima SOB genome, and appears to have arisen from several horizontal transfer events from other symbiont taxa or free living populations (Figure 3B).

**Figure 3.**
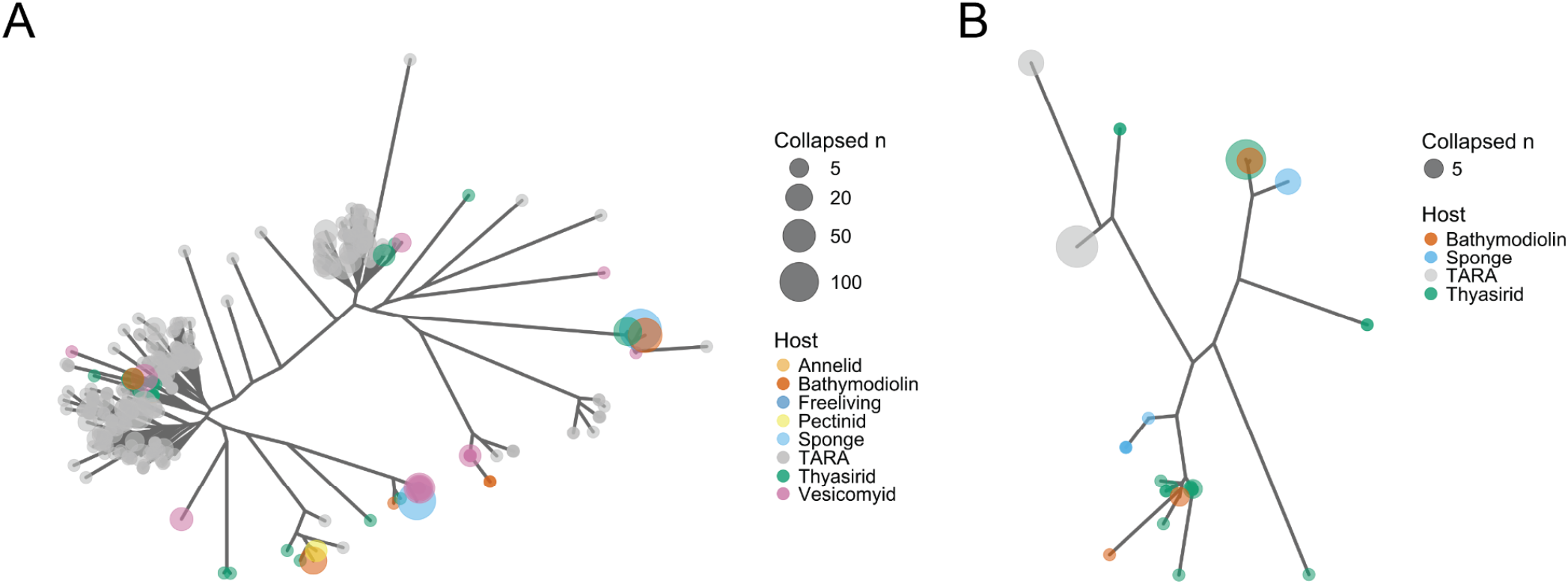
Insertion sequence diversity of *Isorropodon fossajaponica* SOB and *Thyasira* sp. Haima SOB. Unrooted phylogenies of IS sequences and similar BLAST hits from the NCBI nt and TARA Oceans databases for (A) IS3 in *Isorropodon fossajaponica* SOB and *Thyasira* sp. Haima SOB, and (B) IS5 in *Thyasira* sp. Haima SOB.

### IS elements disrupted synteny, mediating structural rearrangements genome-wide

The characteristic syntenic genome structure of vesicomyid symbionts was disrupted in the two lineages with abundant IS element copy numbers. Pairwise whole-genome alignments (Figure 4) showed that synteny was largely conserved among most symbionts, with a limited number of inversions, but was almost entirely lost in the *I. fossajaponica* SOB genome. The *Thyasira* sp. Haima SOB genome also had many inversions relative to the other strains, but global synteny was still apparent in the order of the inverted blocks (blue dot plots in Figure 4). Local homology at IS copies scattered across the genomes can be seen when the genomes are compared to themselves (zoomed-in windows in Figure 4). When scattered around the genome (*e*.*g*., Figure 2C), IS elements may provide sites of local homology for ectopic recombination to occur, inducing structural mutations such as insertions, deletions, and inversions.

**Figure 4.**
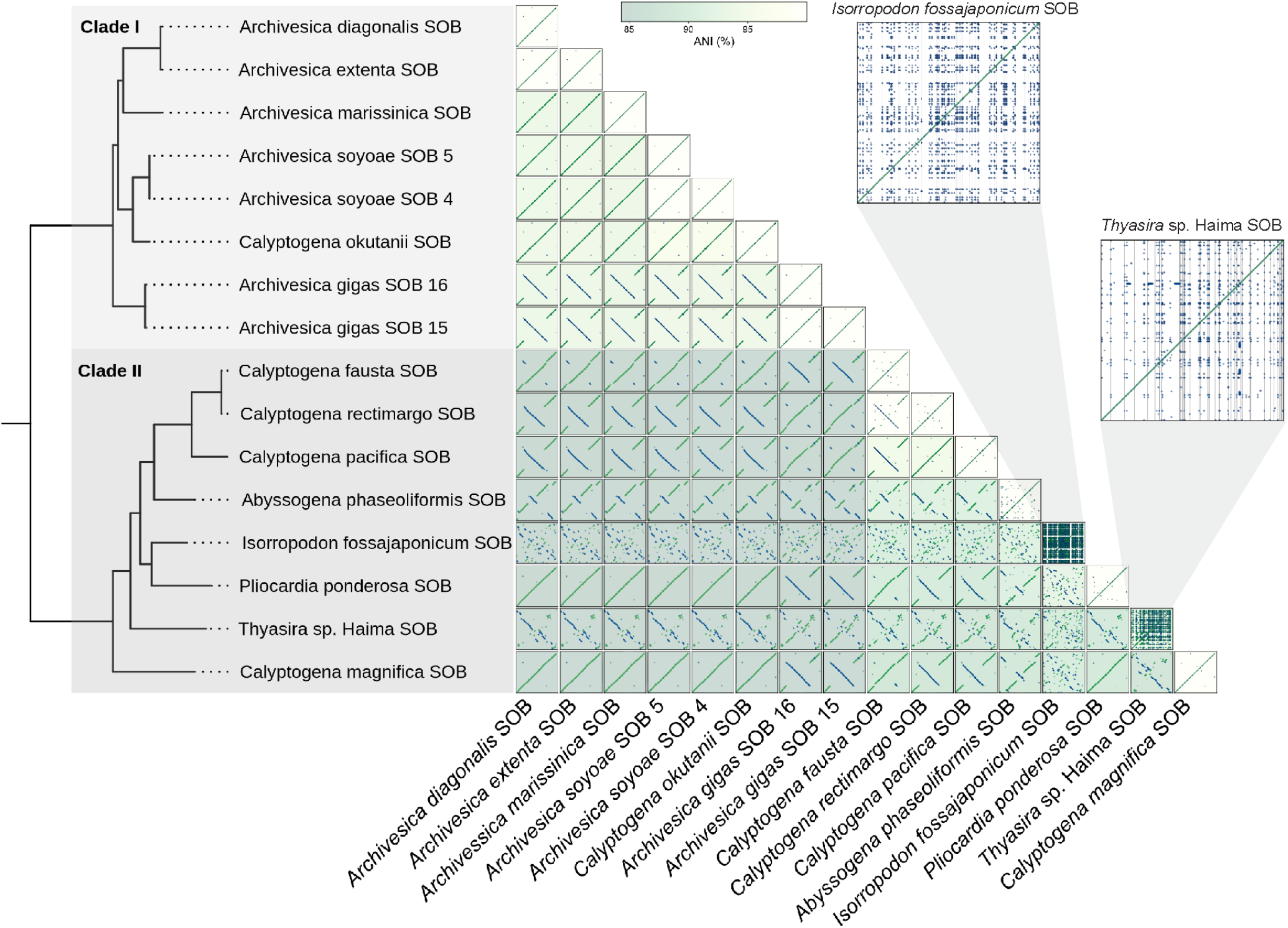
Lineages with phage-derived insertion sequences have highly rearranged genomes. An all-vs-all whole-genome alignment matrix of all Clade I and Clade II symbiont genomes. Green lines indicate alignments in the same direction. Blue lines indicate inversions. Alignment plots are colored by the average nucleotide identity (ANI) across all aligned segments. The self-alignments of *Isorropodon fossajaponica* SOB and *Thyasira* sp. Haima SOB are enlarged to show detail of duplicated elements, with the location of insertion sequences shown as vertical gray lines.

### Clade II vesicomyid symbionts experienced gene gain and duplication events after divergence from an obligate ancestor

Gene gains and duplications occurred more frequently throughout Clade II symbionts than Clade I symbionts, even after excluding predicted IS elements, prophages, and other putative HGT regions. Gene gains, which are classified as an orthogroup in which at least one copy is present in all children of a given node and not in its own ancestor, are enriched in Clade II symbionts (Figure 5A). The *Abyssogena phaseoliformis* SOB has the highest number of predicted gene gain events, at 196 gains. The *Thyasira* sp. Haima genome has a relatively low number of gains predicted compared to other expanded-genome symbionts, and is lower than the non-expanded genome of the *Pliocardia ponderosa* SOB, indicating that excessive gene gain has not resulted from the acquisition of mobile elements. The *Isorropodon fossajaponica* SOB has 127 gene gains, and its sister taxa the *Pliocardia ponderosa* SOB has 98. The relatively high number of gene gains in both the expanded genome containing IS elements and its relatively smaller genome relative could suggest that both genomes have novelly acquired genes via mobile elements not detected in the *Pliocardia ponderosa* SOB. Alternatively, gene gain events could have arisen through alternative mechanisms of gene gain that arise from ancestral coding sequence. One mechanism is the rapid neofunctionalization or frameshift mutations, of ancestral or duplicated genes. Additionally, novel genes can sometimes arise *de novo* from non-coding sequences [28–30]. However, these events are difficult to distinguish from gene loss, which occurs frequently in ancient symbiont lineages, including vesicomyids (Supplementary Figure S3). Considering pseudogenization could also create open-reading frames that are unique from their ancestral orthogroup beyond detectable homology, it is difficult to distinguish these events from events of pseudogenization and gene loss without functional characterization.

**Figure 5.**
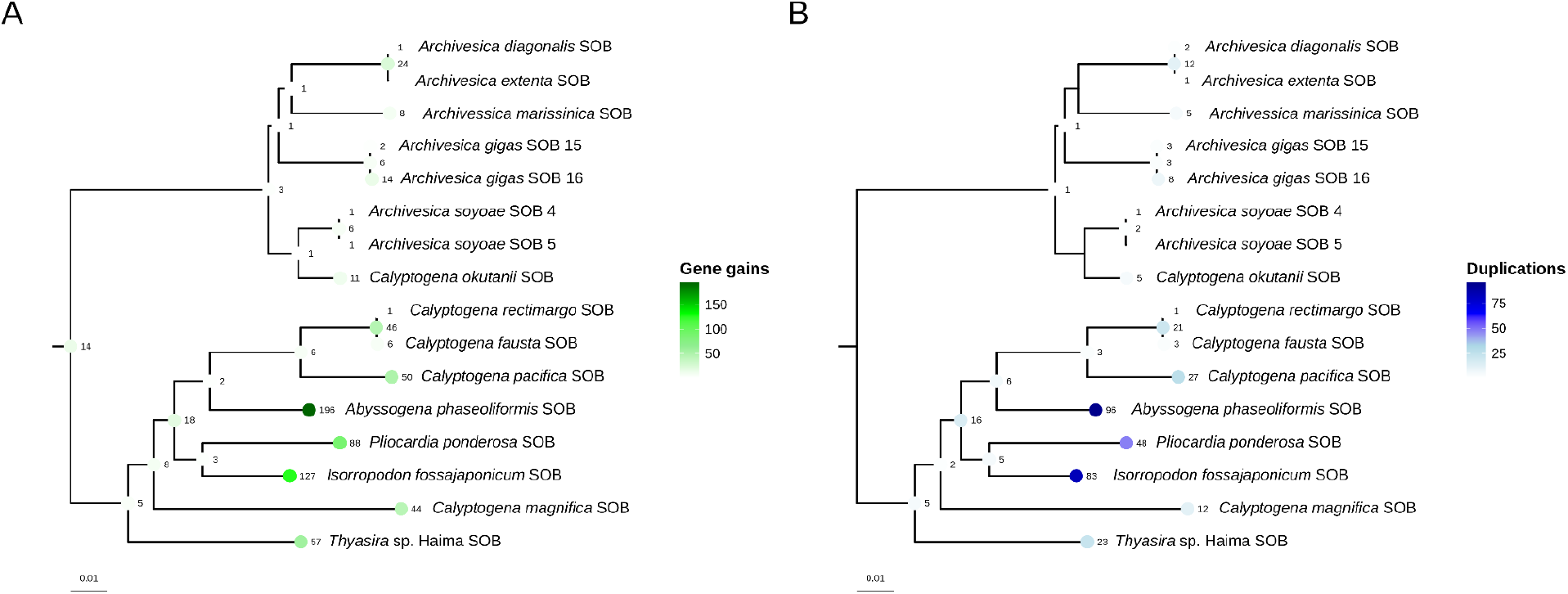
Gene gains and duplications across the phylogeny of Clade I and Clade II vesicomyid symbiont genomes. (A) Gene gains and (B) duplications, excluding IS elements, prophages, and putative HGT regions. The phylogeny was constructed using the STAG algorithm on orthogroup gene trees, output by Orthofinder v. 3.1.0.

Gene duplications, which are classified as a node in an orthogroup gene tree in which at least one genome carries copies in more than one descendant clade, are also enriched in Clade II symbionts (Figure 5B). The *Abyssogena phaseoliformis* SOB has 96 predicted gene duplications, closely followed by 83 gene duplications in the *Isorropodon fossajaponica* SOB. Similarly to gene gains, gene duplications have occurred more frequently in the non-expanded *Pliocardia ponderosa* SOB genome than in the expanded *Thyasira* sp. Haima SOB genome. This implies that selection for gene duplications could fuel adaptation independently within these lineages. Further characterization of the function of these genes is required to better understand if these genes are functional, if they are undergoing neofunctionalization, and whether they provide an adaptive benefit.

### A host-switching symbiont contains novel horizontally-acquired and expanded genes encoding large adhesins

The *Thyasira* sp. Haima SOB harbors multiple copies of an unusually large gene with structural homology to known adhesins, suggesting a specialized role in symbiont attachment or host cell interaction. Four large (∼24,000–40,000bp), highly repetitive genes are present in the Thyasirid symbiont *Thyasira* sp. Haima SOB, which is within Clade II of the vesicomyid symbionts (Figure 6A). Functional domain prediction, structural prediction, and structural homology searches indicate domains with high homology to known adhesins found across highly diverse prokaryotes and eukaryotes, indicating likely adhesin function (Figure 6B). Three of the four large adhesins are highly similar to each other, therefore we refer to these as Group A adhesins, and the fourth adhesin as Group B. Protein BLAST searches of Group A adhesins reveal no hits to known sequences on the amino acid level. On a nucleotide level, BLAST searches reveal hits to a shorter homolog found within the same symbiont genome and shared with other Clade II symbionts. BLAST searches of Group B yield hits with ∼50% identity to distantly related bacteria on the amino acid level, and a single repeat of 72% identity on the nucleotide level to a free-living relative within Thioglobaceae. Insertion sequences flank all on one or both sides, which might have mediated their transposition and expansion during recombination events. Considering *Thyasira* sp. Haima SOB occupies a Thyasirid host, which typically hosts extracellular or horizontally-transmitted symbionts, large adhesins might aid in the invasion of host cells or exchange of metabolites along the surface of host cells.

**Figure 6.**
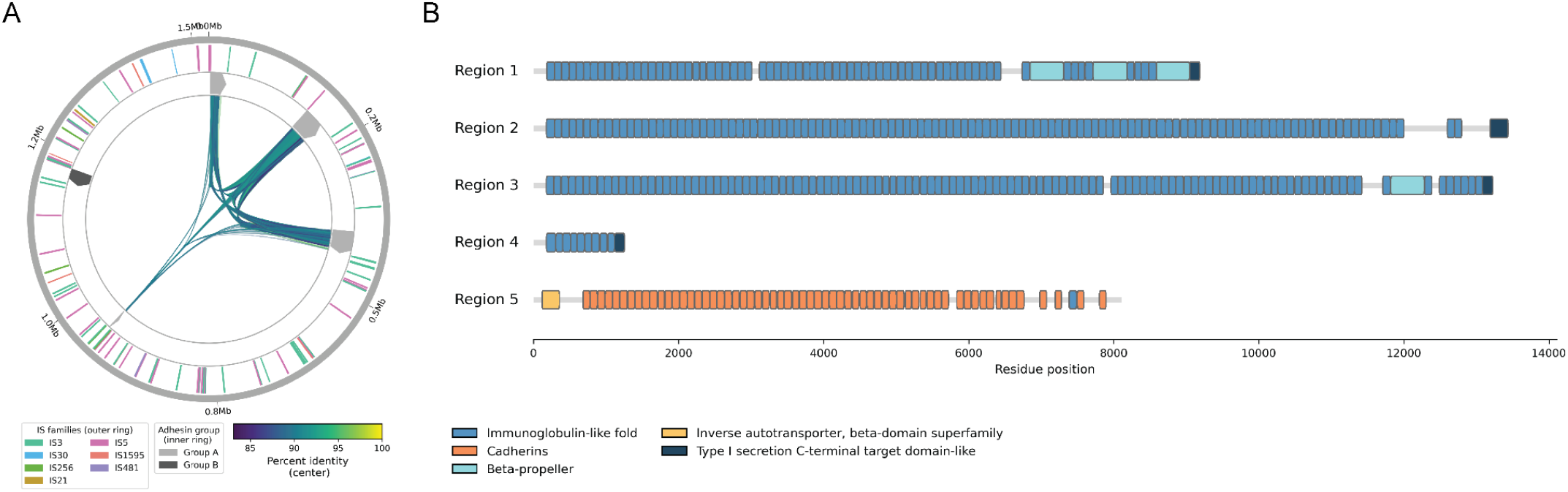
Large genes in the Thyasira sp. Haima SOB genome encode novel adhesins. (A) IS element distribution, colored by IS family (outer ring), distribution of adhesins, colored by group (inner ring), and BLASTn self-alignment hits across the genome, colored by percent identity (center). (B) Predicted functional domains across adhesins.

### Retention of recombination machinery may facilitate genome expansion

The acquisition and subsequent expansion of mobile genetic elements raises the question of which molecular mechanisms enable these events, and whether the retention of recombination machinery contributes to the evolutionary flexibility of these symbiont genomes. Although genes involved in homologous recombination and DNA repair are frequently lost during endosymbiont genome reduction, as observed in Clade I symbionts, they are largely retained across Clade II, consistent with previous analyses [31, 32] (Supplementary Figure S4). The retention of homologous recombination machinery may be particularly relevant to genome expansion, as it could facilitate both the integration of foreign genetic material and its subsequent proliferation through duplication and genomic rearrangement, as suggested by the duplications and repeat expansions of the adhesin-like genes in *Thyasira* sp. Haima SOB. However, evidence of homologous recombination has been detected even in vesicomyid symbionts lacking components of canonical recombination pathways, suggesting that the loss of this machinery does not necessarily preclude recombination or subsequent genome expansion.

Furthermore, because vesicomyid symbionts are transmitted vertically via the surface of host eggs, they may encounter environmental sources of mobile genetic elements during transmission, providing opportunities for horizontal acquisition. The subsequent persistence and fixation of acquired elements may depend on the frequency of their introduction, transmission dynamics, and any selective advantages they confer within the host environment. Nevertheless, the broad retention of canonical recombination and DNA repair machinery across Clade II, despite genome expansion being restricted to only certain lineages, suggests that recombination capacity alone is insufficient to explain these events. Instead, additional factors, potentially arising from differences in symbiont genomic backgrounds, opportunities for horizontal acquisition, or host-associated selective pressures, likely influence whether newly acquired genetic material is retained and subsequently expanded. Further investigation into the activity of retained recombination pathways, the mechanisms of mobile element acquisition, and the fitness consequences of newly acquired genes is necessary to disentangle their respective contributions to genome expansion in these ancient symbiont lineages.

## Conclusion

Our findings challenge the assumption that endosymbiont genome evolution proceeds toward erosion, only stalled by the recovery of previously lost function, but never expanded into new capabilities. By analyzing comparative genomic data across the vesicomyid and vesicomyid-like symbionts, we show that at least three lineages within Clade II have independently expanded their genomes by acquiring novel genetic material via IS elements, phages, and gene duplications. Critically, this new content has not been subject to the same pseudogenization pressures that typically accompany symbiont genome reduction, suggesting it has been retained because it confers a selectable benefit rather than persisting as neutral or nearly-neutral debris awaiting deletion.

The clearest evidence for functional innovation comes from *Thyasira* sp. Haima SOB, whose four large, repetitive adhesin-like genes bear the signatures of both horizontal acquisition and IS-homology mediated duplication. The timing of this gene gain coincides with a shift from a vesicomyid to a thyasirid host, suggesting it may have played a direct role in enabling the host switch by providing an adaptive function. This raises the possibility that mobile genetic elements can act not merely as agents of genomic decay, as is typically emphasized in studies of endosymbiont genome evolution, but as vectors that introduce the specific functional novelty needed to colonize a new ecological niche.

More broadly, these results suggest that the genomic trajectory of intracellular symbionts may be more evolutionarily flexible than the classic model of unidirectional erosion implies. Rather than genome reduction representing an evolutionary dead end, our data raise the possibility that mobile elements provide a recurring mechanism by which eroded genomes can acquire new functional capabilities. This is a capacity that may be especially important for host-switching, where the demands of a new host environment can differ substantially from those of the old. Future work testing whether these adhesin genes are expressed and functionally deployed during host colonization, and whether similar IS-mediated gene gain occurs in other host-switching symbiont lineages, will help establish how general this mechanism of functional innovation may be across intracellular symbioses.

## Methods

### Data acquisition and quality assessment

We downloaded all available genome assemblies of vesicomyid sulfur-oxidizing symbionts and one closely related thyasirid symbiont (genera *Candidatus Vesicomyidisocius, Candidatus Ruthia, and Candidatus Ruthturnera*) from NCBI (accessed July 27, 2026). We downloaded only complete, circular genomes, totaling 16 individual genomes (Table S1). To assess genome completeness, we used BUSCO v. 6.1.0 [33] with the gammaproteobacteria database in OrthoDB v. 12.2 [33].

### Genome annotation, gene clustering, and mobile element identification

We predicted and annotated genomic features using Prokka v. 1.14.6 [34]. We used Orthofinder v. 3.1.0 [35] with default settings to cluster amino acid sequences within and across symbiont genomes into orthogroups, or groups with potential shared ancestry and function.

Since prokka annotations are based on homology to known sequences and may not identify novel or fragmented transposases, we predicted and annotated additional insertion sequences using ISEScan v. 1.7.3 [36]. An IS21 element that was predicted as a single copy in all 16 genomes was removed due to its location within dnaA considering its known homology to ISTB proteins encoded in IS21 [37].

Considering that many sequences, in addition to the insertion sequences, were unannotated due to no identifiable homolog, we assessed the symbiont genomes or the presence of additional potentially novel mobile genetic elements. We identified prophages using PHASTEST v. 1.0.1 [38]. We classified additional putative HGT regions as regions of the genome >15 kb in which no genes were found to have significant BLASTn or BLASTp hits against the NCBI nt and nr databases, respectively, (e-value threshold of 1e-5) to other relatives within Thioglobaceae. We set the boundaries of putative HGT to the gene boundaries of these stretches. We calculated GC content across the genome and GC content at the third codon position (GC3) of each gene using a custom python script. We calculated the codon adaptation index using the CAI package in python [39], using the codon usage of single-copy orthologs across Clade I and Clade II symbiont genomes as the reference sequence set. We fit a linear regression of CAI on GC3 across all genes. To determine if the putative HGT regions fall below this trend more often than due to chance, we assessed both regions independently. For each putative HGT region, we calculated the signed mean residual from the line of all genes within the region. We generated sliding windows of the length of the region and a step of the average genome size across the genome (441 bp), and assigned genes to a window if their midpoint falls within the window. We used the signed mean residuals of genes within each window as the null distribution, and calculated a p-value as the proportion of signed mean residuals at or below the observed value for the given region.

We identified ORFans, or open reading frames (ORFs) belonging only to a single genome, as orthogroups containing genes from only one genome and no significant BLASTp hits (e-value threshold of 1e-5) in the NCBI nr database (accessed April 3, 2026).

### Insertion sequence phylogenetic analysis

We extracted the nucleotide sequences of predicted IS elements and grouped them by predicted family and across all genomes. For each IS family, we ran a BLASTn search against the NCBI nt and TARA Oceans prokaryotic database using an e-value cutoff of 1e-5. We filtered results to remove hits with query coverage less than 50% and identity less than 60%. We extracted the aligned portion of remaining hits and all IS elements and re-aligned their hits using MAFFT v. 7.526 [43]. We inferred the unrooted maximum likelihood (ML) phylogeny for each IS family using IQTree v. 3.0.1 [44] using MFP to run ModelFinder Plus, with 1000 ultrafast bootstrap replicates.

### Phylogenetic analysis of symbiont genomes

We identified the single-copy orthologs present across all genomes output by Orthofinder described in “Genome annotation, gene clustering, and mobile element identification”. We aligned the amino acid sequences of resulting orthologs using MAFFT v. 7.526 [43], then converted to codon-aware nucleotide alignments using PAL2NAL [45]. We inferred the whole-genome symbiont phylogeny using a partitioned ML approach with IQTree v. 3.0.1 [44], using MFP+MERGE to run ModelFinder Plus and allow IQTree to merge partitions with PartitionFinder, and 1000 ultrafast bootstrap replicates.

### Phylogenetic analysis of host genomes

We retrieved the mitochondrial genomes for all hosts from GenBank (Supplementary Table S3). We extracted nucleotide sequences using GenBank annotations for each of the following genes: 12S, 16S, atp6, atp8, cob, cox1, cox2, cox3, nad1, nad2, nad3, nad4, nad5. Only the sequence for cox1 was present for Thyasira sp. Haima, but considering it is a distant relative of the vesicomyids, it was used only to root the tree. We aligned the nucleotide sequences for each gene using MAFFT v. 7.526 [43]. We then inferred the whole-genome symbiont phylogeny using a partitioned ML approach with IQTree v. 3.0.1 [44], using MFP+MERGE to run ModelFinder Plus and allow IQTree to merge partitions with PartitionFinder, and 1000 ultrafast bootstrap replicates.

### Whole genome alignment

We reoriented the symbiont genomes so that dnaA is the starting position on the forward strand using dnaapler v. 1.4.0 [46]. To identify syntenic regions between each genome, we performed an all-versus-all nucleotide alignment between the genomes of all symbionts using mummer v. 4.0.1 [47] with ‘--maxmatch’ to include repetitive matches. We calculated the average nucleotide identity (% ANI) for aligned segments using mummer.

### Pathway completeness and pseudogenization analysis

To group genes into functional pathways, we used eggNOG-mapper v. 2.1.13 [48] to annotate function from the eggNOG database [49], and then used KEGGaNOG v. 1.1.19 [50] to map eggNOG annotations to KEGG pathways. We measured pathway completeness as the proportion of steps in the KEGG pathway in which the genes necessary for function are present.

We identified pseudogenes using the ‘annotate’ function in Pseudofinder v. 1.14.6 [34] with default settings, using all 61 available Thioglobaceae NCBI RefSeq annotations (accessed July 27, 2026) as the input database for homology search. For each symbiont genome analyzed, we removed its own genome from the database, if present, to reduce bias in pseudogene calls due to all genes being present.

### Gene gain, duplication, and loss reconstruction

To quantify gene gain and duplication in vesicomyid symbionts, we reconstructed gene family evolution across the vesicomyid symbionts and their relatives within the Thioglobaceae. By including the less reduced genomes of free-living relatives and horizontally-transmitted bathymodiolin symbionts, we reduce the likelihood that traces of ancestral retention will be called gains. All 37 available complete Thioglobaeae genomes, including vesicomyid and vesicomyid-like symbionts, free-living relatives, and symbionts of bathymodiolus bivalves, were downloaded from the NCBI genome database (accessed July 27, 2026). Genes were predicted using the same method described in “Genome annotation and gene clustering.” Predicted genes within the vesicomyid and vesicomyid-like symbiont genomes were filtered to remove genes that overlap insertion sequences, prophages, or putative HGT regions, predicted above, by more than 50% of the gene length. Orthofinder v. 3.1.0 [35] was used to predict orthologous genes across all 37 genomes. To map gene gains, duplications, and losses to the phylogeny, a modified version of GLADE v. 1.0.0 [51] was used directly on the OrthoFinder output using default settings. Since the expanded genomes may contain duplicates of HGT-acquired genes, which are otherwise counted as duplications despite having an unknown number of copies in its source genome, an additional filter requiring orthogroups to have copies in at least two genomes was added. Additionally, GLADE utilizes gene trees and therefore requires a minimum of 4 total gene copies in an orthogroup, therefore it may severely undercount gene gains that occur at nodes with less than 4 taxa, including leaf nodes. Therefore, a custom python script was used to trace gene gains in orthogroups with less than 4 total copies. Orthogroups which did not have a copy present at every child of the most recent common ancestor were discarded to prevent falsely classifying widespread losses and sparse retention as gains. The resulting gains were added to the output of the modified GLADE output.

### Structural and functional prediction of novel adhesins

To group adhesins and identify other regions of the genome with similar sequence content, a BLASTn search was conducted between the whole *Thyasira* sp. Haima SOB genome and itself. To predict functional domains in group A and group B adhesins in the *Thyasira* sp. Haima SOB genome, we used Interproscan 5.76-107.0 [52]. We extracted unique domains and predicted their structures using AlphaFold 3 [53]. To search for proteins with similar predicted structure without limiting to high amino acid sequence similarity, we used the predicted structures as input to Foldseek [54].

## Data availability

Scripts and commands used for analysis can be found at https://github.com/moreyc/symbiont_genome_de-erosion.

## Acknowledgements

This work was supported by UC Santa Cruz and the NIH: R00GM135583 and R35GM157189 to SLR.

