## Supplementary Figures for "Accretion of novel genetic elements reverses genome erosion and enables host-switching in the deep sea"

Supplemental Figures

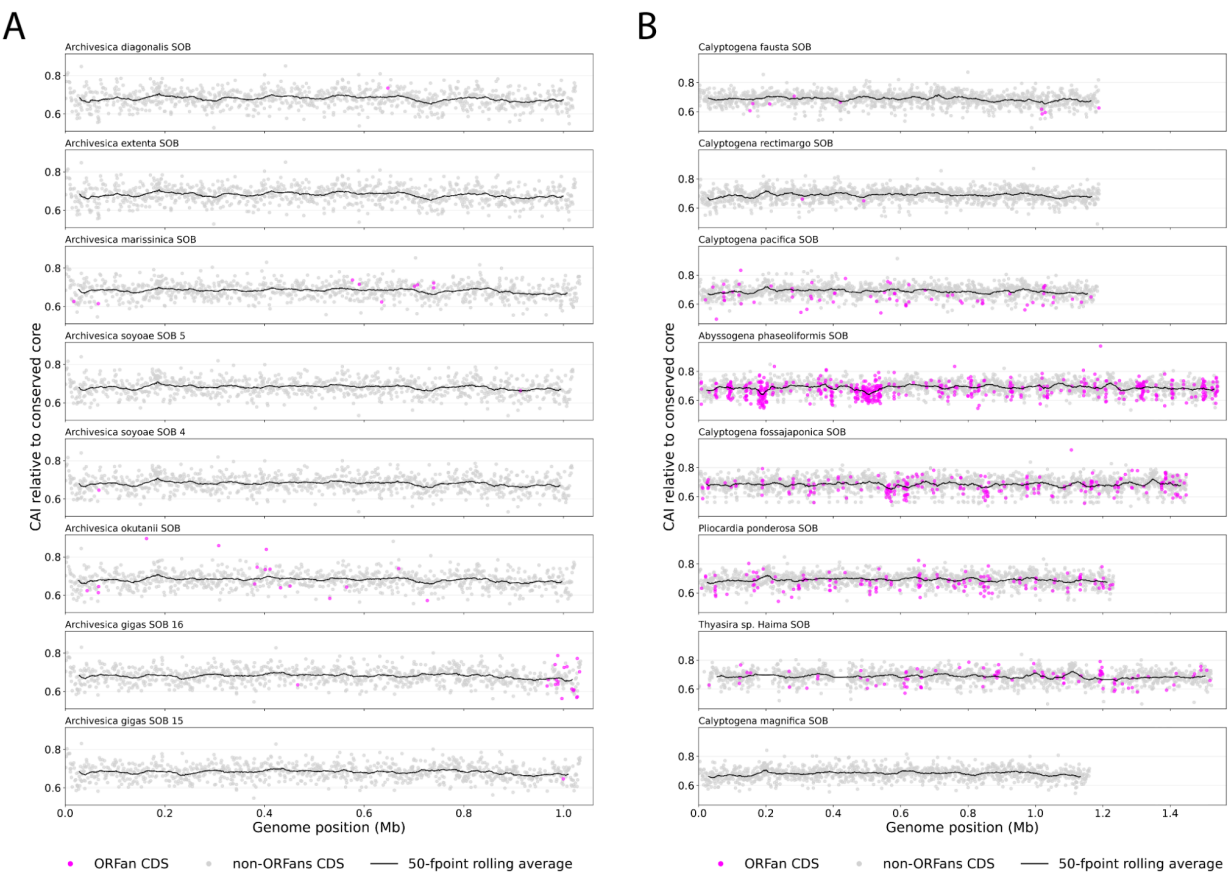

**Figure S1.** Codon usage bias and ORFan distribution across vesicomyid and vesicomyid-like symbiont genomes for (A) Clade I and (B) Clade II.

**Figure S2.** Phylogenetic trees of insertion sequences and their closest hits in BLAST and TARA Oceans databases. IS elements are clustered into trees based on predicted IS family.

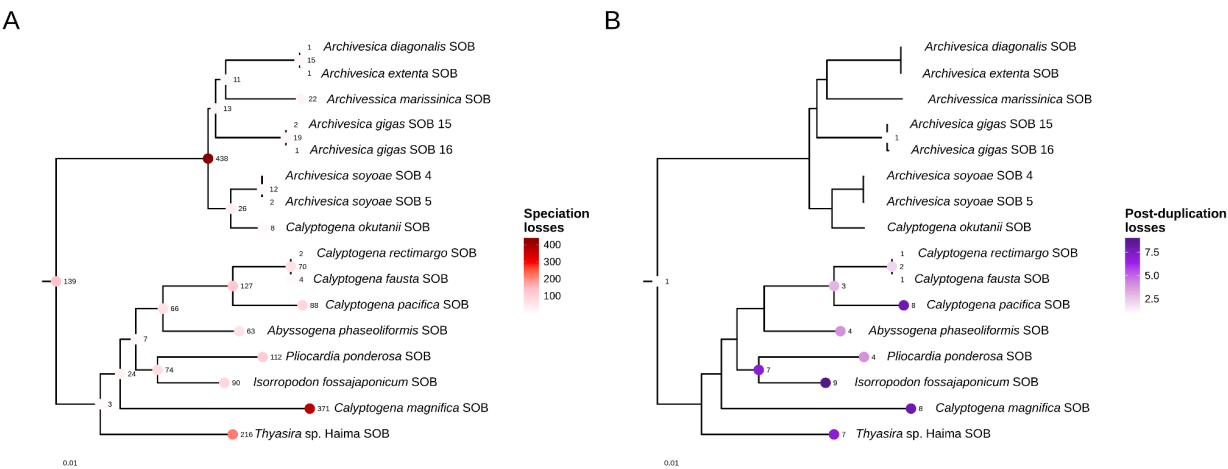

**Figure S3.** Losses of genes and duplicated genes across the phylogeny of Clade I and Clade II vesicomyid symbiont genomes. (A) Speciation losses, or gene losses of non-duplicated copies, and (B) post-duplication

losses, or losses of gene copies that were previously duplicated from an ancestral gene. Gene losses were predicted excluding IS elements, prophages, and putative HGT regions. The phylogeny was constructed using the STAG algorithm on orthogroup gene trees, output by Orthofinder v. 3.1.0.

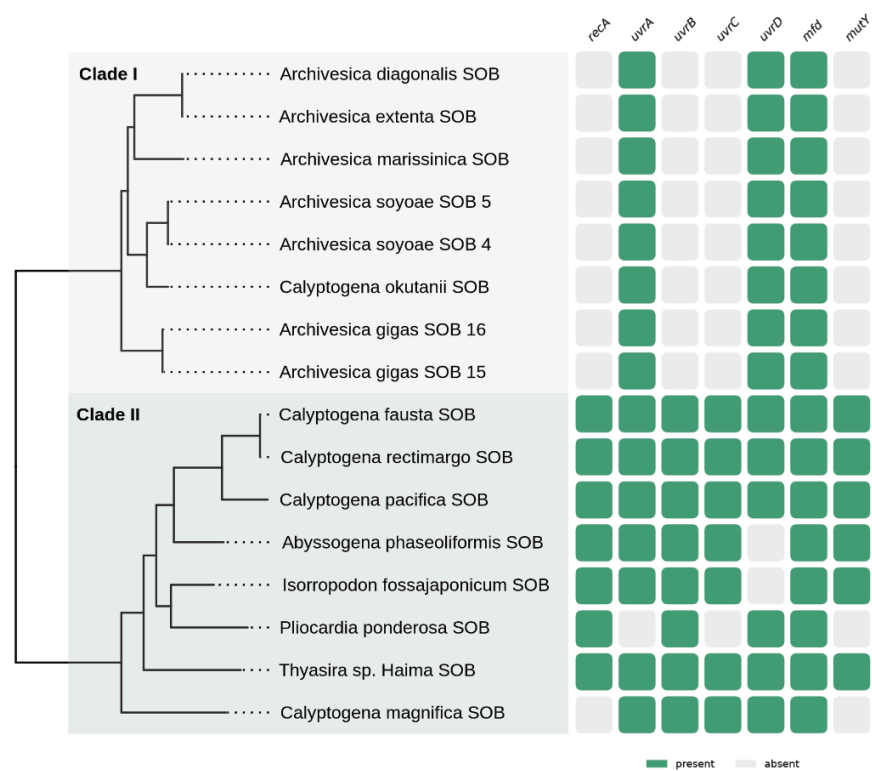

**Figure S4.** Presence or absence of primary recombinase *recA* and DNA-repair genes across vesicomyid and vesicomyid-like symbionts.

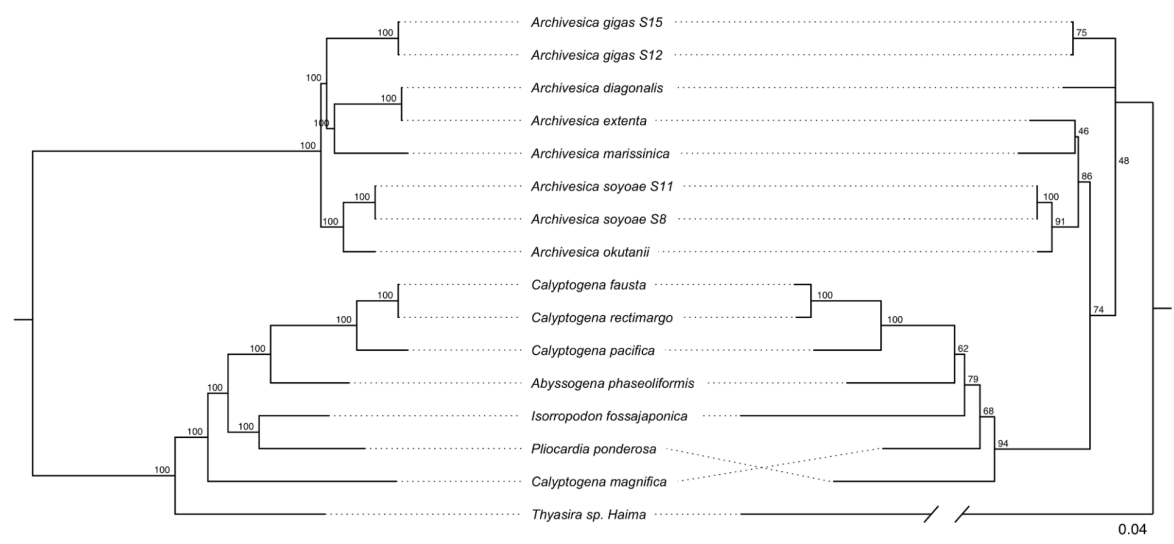

**Figure S5.** Host and symbiont phylogenetic congruence. Maximum-likelihood phylogenies of (Left) symbionts, using single-copy orthologs, and (Right) hosts, using 13 mitochondrial genes.
